# Benchmarking functional connectivity by the structure and geometry of the human brain

**DOI:** 10.1101/2021.10.12.464110

**Authors:** Zhen-Qi Liu, Richard F. Betzel, Bratislav Misic

**Affiliations:** McConnell Brain Imaging Centre, Montréal Neurological Institute, McGill University, Montréal, Canada; Psychological and Brain Sciences, Indiana University, Bloomington, IN, USA

## Abstract

The brain’s structural connectivity supports the propagation of electrical impulses, manifesting as patterns of co-activation, termed functional connectivity. Functional connectivity emerges from the underlying sparse structural connections, particularly through poly-synaptic communication. As a result, functional connections between brain regions without direct structural links are numerous, but their organization is not completely understood. Here we investigate the organization of functional connections without direct structural links. We develop a simple, data-driven method to benchmark functional connections with respect to their underlying structural and geometric embedding. We then use this method to re-weigh and re-express functional connectivity. We find evidence of unexpectedly strong functional connectivity within the canonical intrinsic networks of the brain. We also find unexpectedly strong functional connectivity at the apex of the unimodal-transmodal hierarchy. Our results suggest that both phenomena – functional modules and functional hierarchies – emerge from functional interactions that transcend the underlying structure and geometry. These findings also potentially explain recent reports that structural and functional connectivity gradually diverge in transmodal cortex. Collectively, we show how structural connectivity and geometry can be used as a natural frame of reference with which to study functional connectivity patterns in the brain.

## INTRODUCTION

Axonal wiring among neurons and neuronal populations promotes signal exchange and information integration. At the mesoscale, signaling via the complex network of anatomical projections manifests as patterns of temporal correlations, termed functional connectivity (FC). Functional connectivity is highly organized [5, 13, 47], reproducible [18, 33] and related to individual differences in behaviour [31, 42].

Most pairwise functional connections are not supported by a direct structural connection. By definition, functional networks are fully connected, while structural networks are sparse (Fig. 1). Across species, reconstruction techniques and spatial scales, structural connection density is typically reported to be between 2% and 40% [48] (but see also [26]), meaning that the majority of functional connections between two regions are not accompanied by a corresponding direct structural connection. These “indirect” functional connections are thought to emerge from polysynaptic communication in the structural network [4, 45].

**Figure 1.**
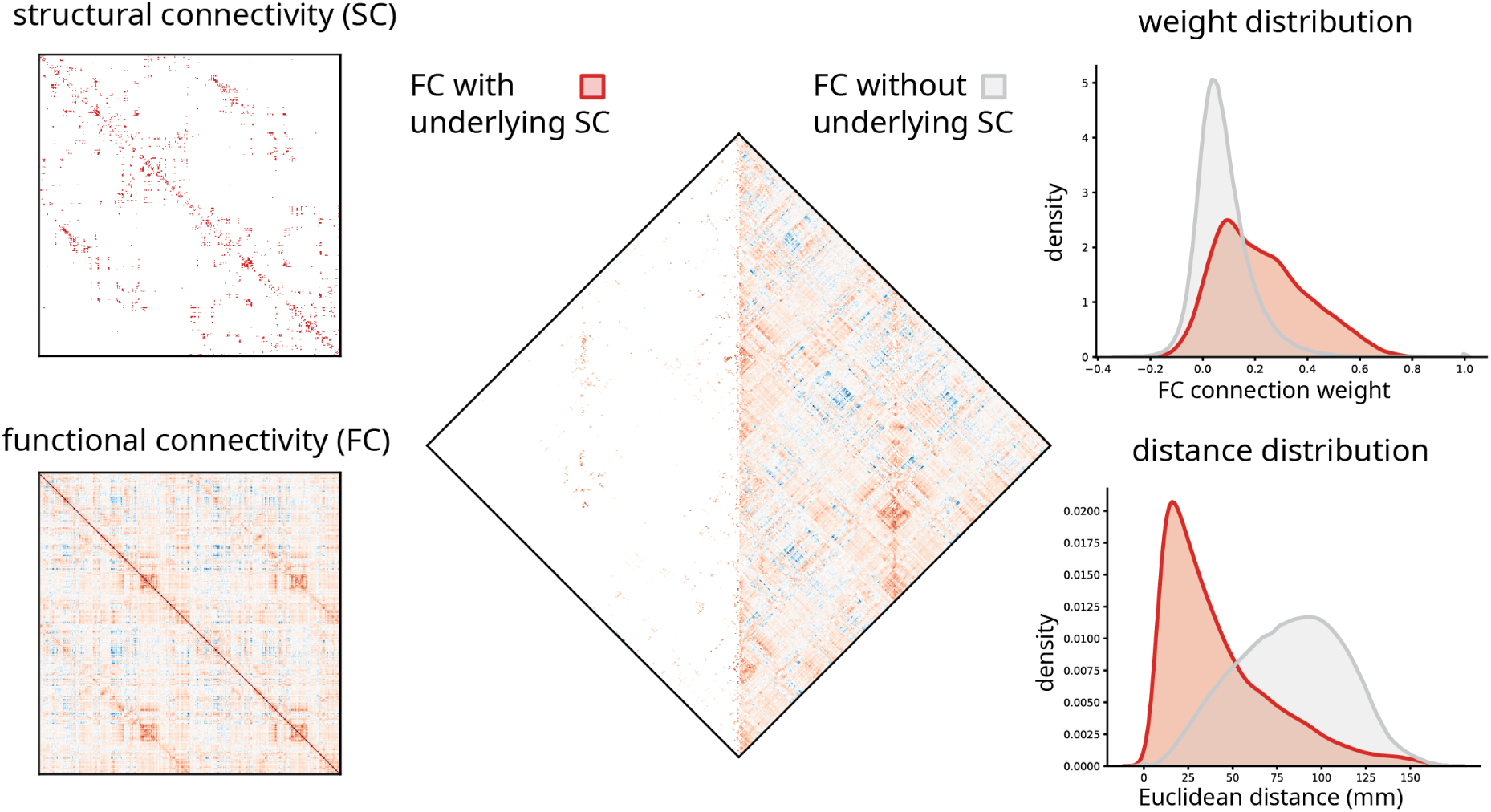
Functional connections with and without direct structural links. Left: structural connectivity (SC) and functional connectivity (FC) matrices in the 1000-node Lausanne parcellation [10]. Middle: Functional connections with and without underlying structural connections. Right: The weight and anatomical (Euclidean) distance distribution of the two types of functional connections.

Importantly, structural and functional connectivity are fundamentally constrained by the spatial embedding of brain regions [44]. Structural connection probability is inversely correlated with spatial separation, such that proxmimal neural elements are more likely to be structurally connected, while distant neural elements are less likely to be connected [21, 26, 37]. A similar distance dependence is also observed for functional connectivity [24, 30, 39, 41]. The over-representation of low-cost, short-range connections is thought to reflect finite material and metabolic resources (Fig. 1) [8]. Altogether, structural connectivity and spatial proximity constitute a natural frame of reference for understanding and interpreting functional connectivity.

Here we investigate the organization of functional connections without direct structural links (Fig. 1). We develop a simple method that uses robust relationships between geometry, structure and function as the baseline to re-weigh and re-express functional connectivity. We use the method to identify functional connections that are greater than expected given their structural and geometric embedding. We then show that the arrangement of these connections systematically follows the functional modules (intrinsic networks) [47] and the putative unimodal-transmodal hierarchy of the brain [24].

## RESULTS

The results are organized as follows. We first establish a method to quantify how unexpectedly strong a functional connection is given the physical Euclidean distance between its connected areas. We then describe the organizational principles of these structurally-unconnected functional connections by characterizing their (1) statistical properties, (2) correspondence with intrinsic networks, and (3) correspondence with cortical hierarchies. Data sources include (see *Materials and Methods* for detailed procedures):

- *Structural connectivity*. Structural and functional connectivity were derived from *N* = 66 healthy control participants (source: Lausanne University Hospital; https://doi.org/10.5281/zenodo.2872624) using the 1000-node Lausanne parcellation [10]. Participants were randomly divided into a *Discovery* and *Validation* cohort (*N* = 33 each). Structural connectivity was reconstructed using diffusion spectrum imaging and deterministic streamline tractography. A consistency- and length-based procedure was then used to assemble a group-representative structural connectivity matrix [7, 28, 29].
- *Functional connectivity*. Functional connectivity was estimated in the same individuals using resting-state functional MRI (rs-fMRI). A functional connectivity matrix was constructed using pairwise Pearson correlations among regional time courses. A group-average functional connectivity matrix was then estimated as the mean connectivity of pair-wise connections across individuals.

### Long-range functional connections are unexpectedly strong

To quantify how unexpectedly strong a functional connection is, we first seek to establish a baseline. Fig. 2a shows the relationship between the spatial separation of two nodes (abscissa) and the functional connectivity between them (ordinate). Functional connections that are supported by an underlying structural connection are shown in red, and all other functional connections, which we refer to as indirect or structurally-unconnected FCs, are shown in grey. We note the classical exponential decrease in magnitude with increasing spatial separation [38, 44]. We also note that connected (monosynaptic) and unconnected (polysynaptic) FCs have similar distributions at short distances, but that they diverge considerably at long distances. Namely, when the spatial separation between two regions is greater than approximately 125 mm, there is greater variability among unconnected FCs, with many unconnected FCs marked by greater magnitude than connected FCs spanning comparable distances.

**Figure 2.**
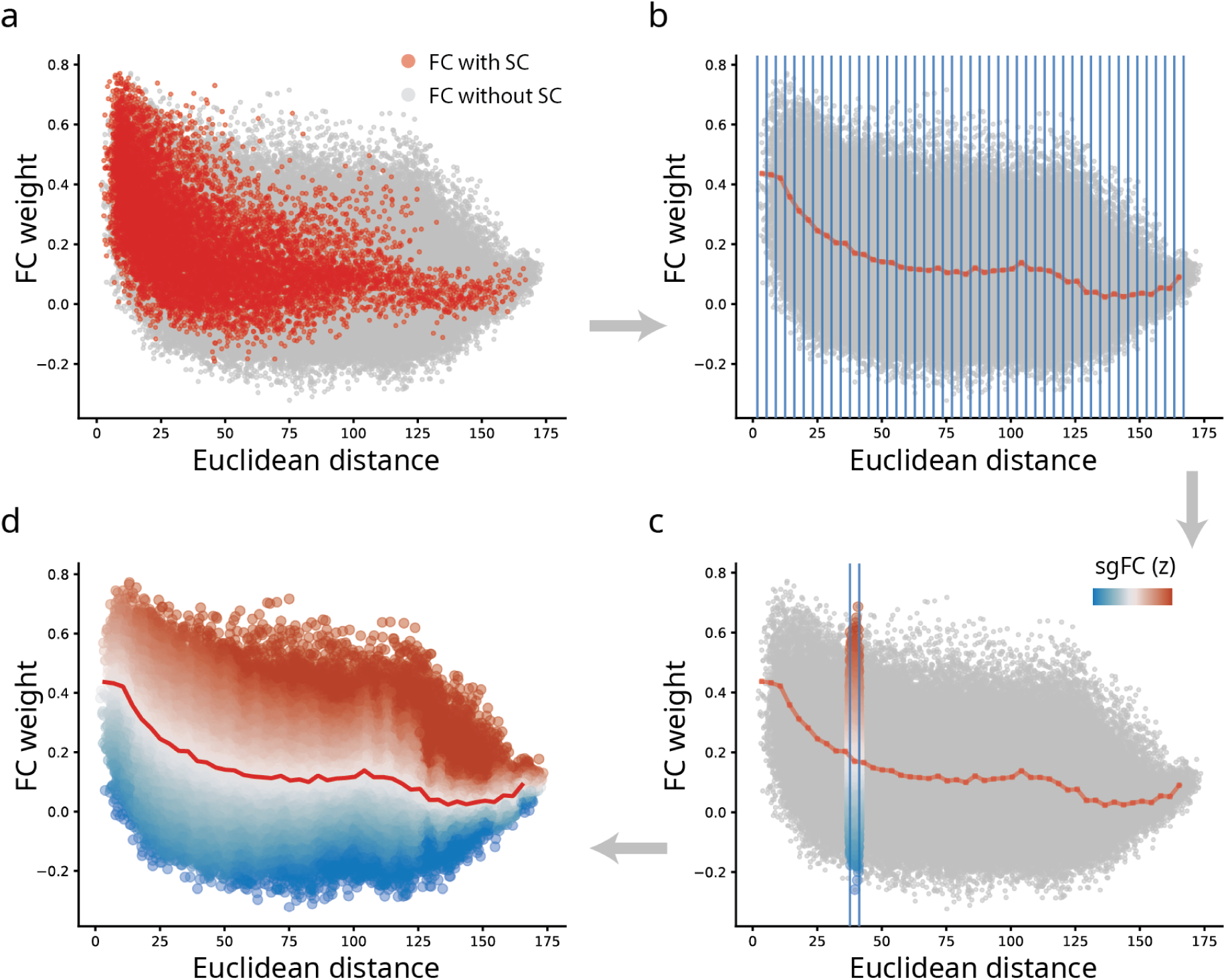
Benchmarking FC by structure and geometry. (a) FC connection weight-to-distance relationship shown for FC with (red) and without (gray) direct SC connections, respectively. (b) FCs grouped into distances bins (blue lines), and the mean value within each bin of those with direct SCs (dotted red line). (c) Within a sample bin, unconnected (polysynaptic) FCs are expressed as a z-score relative to connected (monosynaptic) FCs. We refer to this z-score as structure- and geometry-informed FC (sgFC). (d) sgFCs shown as a smoothly-transitioning spectrum after the procedure is applied for each distance bin. See *Methods* for more technical details and Fig. S1 for details about smoothing and bin size selection.

We therefore set the magnitude of connected FCs at a given distance as the baseline for unconnected FCs at a comparable distance. The goal is to identify unconnected FC that are unexpectedly large relative to connected FCs. To operationalize this intuition, we first bin FCs according to their spatial proximity (Fig. 2b). Within each bin, we record the distribution of connected FCs, including their mean and standard deviation. Finally, we express each unconnected FC as a z-score relative to the distribution of connected FCs in the same distance bin (Fig. 2c). This measure reflects how unexpectedly strong a functional connection is, given its length. Importantly, z-scores for unconnected FCs are estimated based on moments of a distribution estimated for connected FCs. For simplicity, we term the re-expressed unconnected FCs as structure- and geometry-informed FC (sgFC).

Fig. 2d shows the re-weighing of unconnected FCs. Across the entire range of distances, there exist many unconnected FCs that are disproportionately strong relative to their length. A population of unconnected positive FCs spanning distances greater than 125 mm are particularly prominent, suggesting the existence of multiple strong functional interactions above and beyond what would be expected on the basis of their length. In the following sections we explore the organization of these connections in greater detail. For sensitivity analyses regarding bin sizes, preprocessing choices and validation, please see *Control analyses* and Figs. S1,S2. For replication in individual participants, please see Fig. S3

### Contribution to intrinsic network architecture

We first ask how conventional FC and sgFC are related to each other, and how they are distributed within and between intrinsic functional networks [47]. Fig. 3a shows the correlation between FC and sgFC connection weights. As expected, the re-weighing of FCs accentuates some connections and attenuates others.

**Figure 3.**
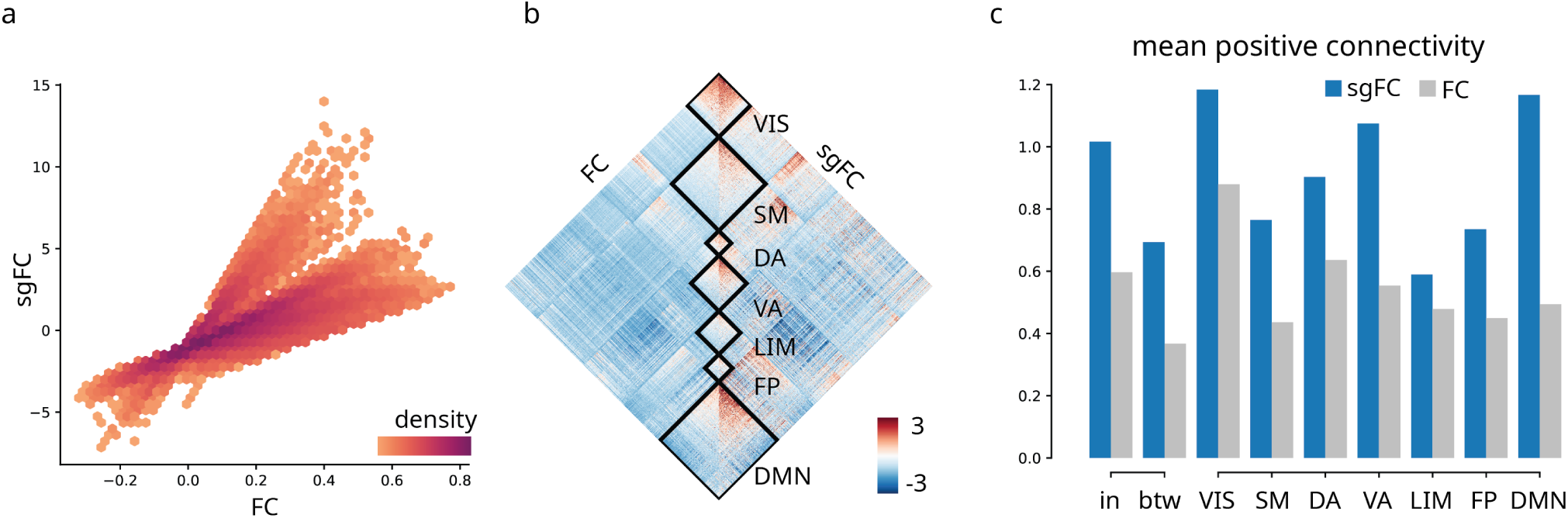
Contribution to intrinsic network architecture. (a) sgFC correlated with FC, colored by scatter density. Only polysynaptic FCs are shown. (b) FC and sgFC shown side-by-side, re-ordered according intrinsic networks [47]. VIS = visual, SM = somatomotor, DA = dorsal attention, VA = ventral attention, LIM = limbic, FP = frontoparietal, DMN = default mode. (c) Comparison of within- and between-network mean positive-valued connectivity with a dissection of within-network connectivity for intrinsic networks. In panels b and c, polysynaptic FCs are standarized by the overall average and standard deviation of FCs with direct SCs to facilitate comparison.

To investigate whether the re-weighing of FCs reflects any organizational features of the brain, we first display FC and sgFC, now re-ordered by the canonical intrinsic networks (Fig. 3b) [47]. The figure shows prominent weights for uncorrected FCs within the diagonal blocks, suggesting that the re-weighing emphasizes within-network connections. Fig. 3c confirms this intuition, showing that re-weighing makes within-network connectivity more prominent. In other words, the well-studied community structure (modules) of functional networks appears to be supported by FCs that are unexpectedly strong given their length.

Interestingly, the largest differences between uncorrected and corrected FCs are observed within transmodal networks (default mode and ventral attention), while more modest differences are observed in the unimodal networks (visual and somatomotor) (Fig. 3c).

This suggests that unexpectedly strong FCs may occur more frequently between brain regions at the apex of the unimodal-transmodal cortical hierarchy. We investigate this possibility in the next section.

### Contribution to the cortical hierarchy

We next investigate the arrangement of unconnected FCs in macroscale cortical hierarchies. Recent work suggests that the functional architecture of human brain networks can be summarized by a small number of smooth topographic gradients, with the most prominent such gradient spanning unimodal to transmodal cortex [24]. This putative hierarchy is thought to support a sensory-fugal representational hierarchy [27], and correlates with spatial variation in cytoarchitecture [34], myelination [22] and gene expression [9].

To place each cortical node along this putative hierarchy, we adapted the diffusion embedding method described by Margulies and colleagues [11, 24, 50] (see *Materials and Methods* for more detail). Fig. 4a shows the topography of the first gradient, differentiating primary sensory and transmodal cortices, replicating the original report [24].

**Figure 4.**
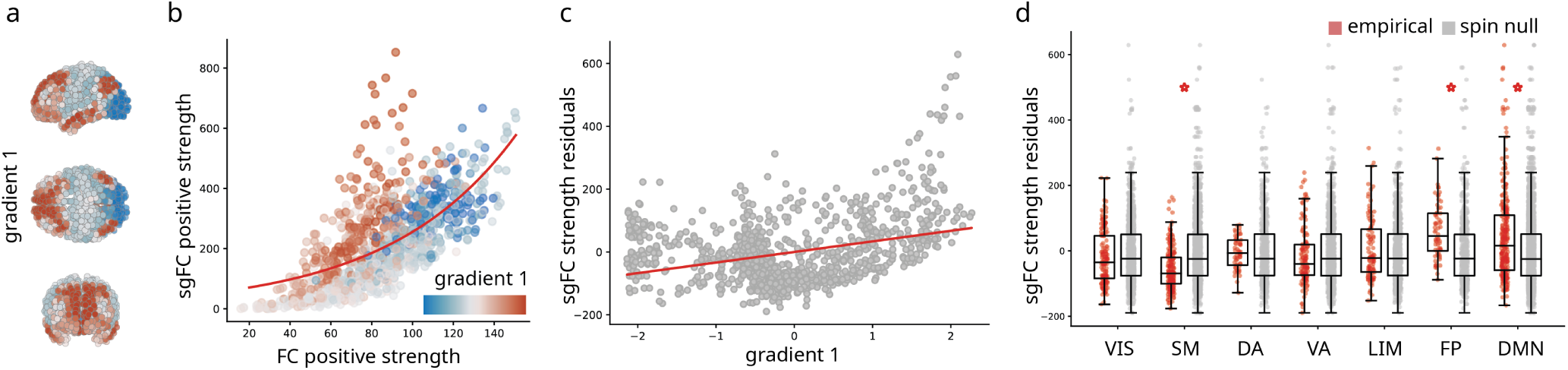
Contribution to the cortical hierarchy. (a) First principal connectivity gradient estimated using diffusion map embedding applied to the FC matrix [24]. Cold colors indicate unimodal regions and warmer colors indicate transmodal regions. (b) Correlation of positive strengths (sum of all weights incident on a given node) between sgFC and FC. Points are regions and are colored by their position in the unimodal-transmodal gradient. An exponential curve is fitted to the points (red line). (c) Residuals of the fitted curve in (b) correlated with gradient 1. (d) Residuals grouped by intrinsic networks and benchmarked against spatial autocorrelation-preserving null models [1, 25]. Statistically significant differences (with Bonferroni correction) are marked with a red asterisk.

To assess the hypothesis that unexpectedly strong FCs are more concentrated in transmodal cortex, we first compare node strengths (the sum of all weights incident on a given region) computed using FC and sgFC. Fig. 4b shows the relationship between node strength for the original FC matrix and for the sgFC matrix. Nodes are coloured by their position in the hierarchy (gradient 1; red = transmodal, blue = unimodal). The relationship is well-fit by an exponential function (*y* = *e*^*x*^; *R*^2^ = 0.44). Importantly, a cloud of red points are consistent outliers, residing above the curve. In other words, brain regions at the apex of the hierarchy are more likely to participate in unexpectedly strong functional interactions.

We further confirm the link between the cortical hierarchy and unexpectedly strong FCs by computing the residual of each node relative to the exponential trend shown in Fig. 4c (Pearson’s *r* = 0.34). Large positive residuals indicate that the node is disproportionately central in the sgFC functional network. Mean residuals for each intrinsic network, ordered by the unimodal-transmodal hierarchy, are shown in Fig. 4d. The greatest increases appear in the frontoparietal (*T* = 5.96, *p* = 1.26 × 10^−7^, *d* = 0.62) and default mode networks (*T* = 5.45, *p* = 1.13 × 10^−7^, *d* = 0.42), when compared to a null model that permutes region labels while preserving their spatial autocorrelation [1, 25]. Collectively, these results show that transmodal cortex participates in polysynaptic FCs that are stronger than expected given their length.

### Control analyses

The results presented in the preceding subsections are potentially contingent on a number of methodological choices, which we explore in detail here. We first replicate the major findings — the distribution of sgFC weights and their involvement in cortical hierarchies — in a validation cohort constructed from *N* = 33 participants. Fig. 5 shows the Pearson correlation of the two results in the *Discovery* and *Validation* cohorts (see Fig. S2 for reproduced result figures). The correlation coefficients for both measures are greater than 0.8 in all cases.

**Figure 5.**
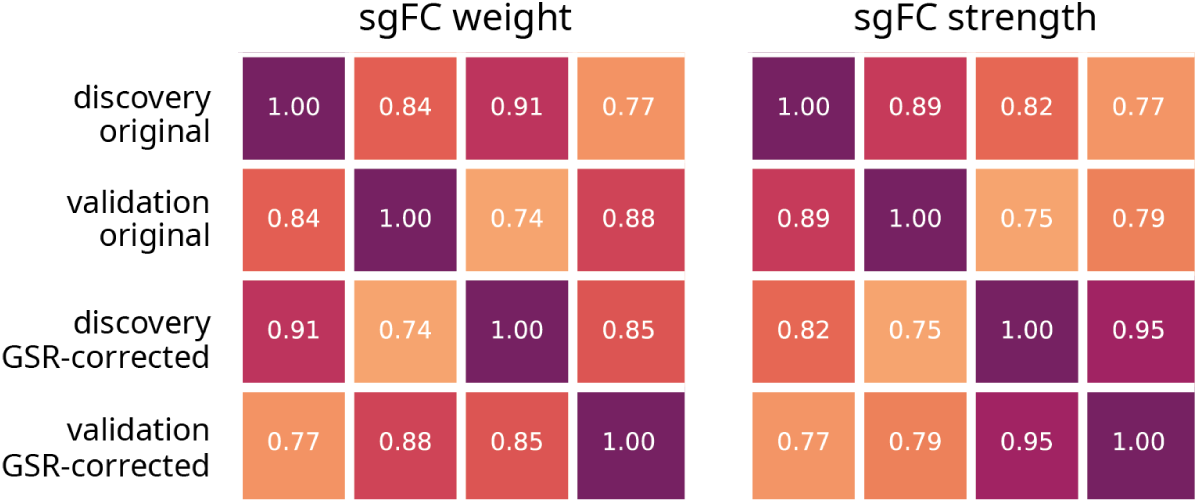
Validation and global signal removal. Correlation matrices shown for sgFC weight (values in Fig. 2d) and sgFC positive node strength (values in Fig. 4b) between controls. Values of sgFC are calculated from *Discovery* and *Validation* datasets, with and without global signal regression (GSR). Reproduced Fig. 2d and Fig. 4b with these values are shown in Fig. S2.

We next seek to determine the extent to which global signal regression could influence the findings. This particular preprocessing step induces negative correlations in FC, profoundly changing the distribution of weights [2, 32]. We re-generated regional time series, correcting for fluctuations in the global signal, and repeated the analysis. Fig. 5 shows the effects of the procedure, in both the *Discovery* and *Validation* cohorts (Fig. S2). As before, there appears to be minimal change in the results, with correlations at approximately 0.9 (for weights) and 0.8 (for strength). In addition, correlations between data cohorts with different processing (e.g. *Discovery* set with no global signal regression correlated with *Validation* set with global signal regression) were also greater than 0.75.

## DISCUSSION

In the present report we introduce a simple data-driven method to benchmark functional connections with respect to their underlying structural and geometric embedding. We find evidence for unexpectedly strong functional connectivity within the canonical intrinsic networks and among transmodal brain regions. These results suggest a hidden but highly organized pattern among polysnaptic FCs.

Our findings build on an emerging literature about the importance of geometry and structural connectivity for functional connectivity in the brain. Although the effect of spatial proximity on the probability and weight of connections is well-known [21, 37], in practice it is less obvious how this information should be taken into account when representing functional connectivity. Likewise, multiple studies report significant correlations between structural and functional connectivity between regions that share direct structural links [20], but how polysynaptic or multi-hop structural connectivity shapes functional connectivity is less well known. Indeed, computational models of structure-function coupling tend to perform more poorly when predicting functional connections between regions that are not structurally connected [17]. More recent communication models of structure-function coupling explicitly account for polysynaptic communication [40, 51]. Here we show that information about structural connectivity and spatial proximity can be naturally used as a frame of reference to describe functional connectivity between regions without direct structural connections.

Interestingly, we find that unexpectedly strong FCs are highly organized with respect to the modular [43] and hierarchical [22] organization of the brain. Although both modules and hierarchies or “gradients” are robust and well-studied features of functional networks, their anatomical origin is less clear [45]. Our results suggest that both phenomena emerge from functional interactions or co-activations that transcend the underlying structure and geometry. In other words, this class of polysynaptic functional connections may be physiologically unique, and future empirical and theoretical studies could potential stratify direct and indirect FCs prior to analysis.

The fact that unexpectedly strong FCs are over-represented in transmodal cortex may potentially explain recent reports that structure-function relationships are regionally heterogeneous. Namely, multiple reports have found that structure-function coupling is greater in unimodal cortex and smaller in transmodal cortex [3, 4, 16, 19, 36, 50, 52]. Our results suggest that the reason for this heterogeneity is that regions in transmodal cortex tend to participate in polysynaptic functional connections that are much stronger than expected given the underlying anatomical constraints. As a result, models relating structural and functional connectivity may be disadvantaged when applied to transmodal cortex relative to unimodal cortex.

In summary, we show how fundamental structural and geometric priors can be used to re-weigh and rerepresent the functional connectivity matrix. Our results show that the canonical features of functional connectivity – modules and hierarchies – are delineated by unexpectedly strong functional connections between nodes without underlying structural links. The biological origin of this class of connections remains an exciting question for future research.

## MATERIALS AND METHODS

### Data acquisition

A total of *N* = 66 healthy young adults (16 females, 25.3 *±* 4.9 years old) were scanned at the Department of Radiology, University Hospital Center and University of Lausanne. The scans were performed in 3-Tesla MRI scanner (Trio, Siemens Medical, Germany) using a 32-channel head-coil. The protocol included (1) a magnetization-prepared rapid acquisition gradient echo (MPRAGE) sequence sensitive to white/gray matter contrast (1 mm in-plane resolution, 1.2 mm slice thickness), (2) a diffusion spectrum imaging (DSI) sequence (128 diffusion-weighted volumes and a single b0 volume, maximum b-value 8000 s/mm^2^, 2.2 × 2.2 × 3.0 mm voxel size), and (3) a gradient echo EPI sequence sensitive to BOLD contrast (3.3 mm in-plane resolution and slice thickness with a 0.3 mm gap, TR 1920 ms, resulting in 280 images per participant). Participants were not subject to any overt task demands during the fMRI scan.

### Structural network reconstruction

Grey matter was parcellated into 68 cortical nodes according to the Desikan-Killiany atlas [15]. These regions of interest were then further divided into four additional, increasingly finer-grained resolutions, comprising 114, 219, 448 and 1000 approximately equally-sized nodes [10]. Structural connectivity was estimated for individual participants using deterministic streamline tractography. The procedure was implemented in the Connectome Mapping Toolkit [12], initiating 32 streamline propagations per diffusion direction for each white matter voxel. To mitigate concerns about inconsistencies in reconstruction of individual participant connectomes [23, 46], as well as the sensitive dependence of network measures on false positives and false negatives [53], we adopted a group-consensus approach [7, 14, 38]. In constructing a consensus adjacency matrix, we sought to preserve (a) the density and (b) the edge length distribution of the individual participants matrices [6, 7, 29]. We first collated the extant edges in the individual participant matrices and binned them according to length. The number of bins was determined heuristically, as the square root of the mean binary density across participants. The most frequently occurring edges were then selected for each bin. If the mean number of edges across participants in a particular bin is equal to *k*, we selected the *k* edges of that length that occur most frequently across participants. To ensure that inter-hemispheric edges are not under-represented, we carried out this procedure separately for inter- and intra-hemispheric edges. The binary density for the final whole-brain matrix was around 2.1%.

### Functional network reconstruction

Functional MRI data were pre-processed using procedures designed to facilitate subsequent network exploration [35]. FMRI volumes were corrected for physiological variables, including regression of white matter, cerebrospinal fluid, as well as motion (three translations and three rotations, estimated by rigid body co-registration). BOLD time series were then subjected to a lowpass filter (temporal Gaussian filter with full width half maximum equal to 1.92 s). The first four time points were excluded from subsequent analysis to allow the time series to stabilize. Motion “scrubbing” was performed as described by Power and colleagues [35]. The data were parcellated according to the same atlas used for structural networks [10]. Individual functional connectivity matrices were defined as zero-lag Pearson correlation among the fMRI BOLD time series. A group-consensus functional connectivity matrix was estimated as the mean connectivity of pair-wise connections across individuals.

### Structure- and geometry-informed indirect FC modelling

To construct the structure- and geometry-informed FC (sgFC), we apply equally-spaced bins to the dimension of Euclidean distance. In each bin, we acquire the mean and standard deviation of those FCs with direct SC link. Then we take the z-score of FCs without direct SC link using the acquired statistics. The final z-scores are smoothed to get a robust representation by averaging over a spectrum of bin numbers (*±*25%) centering the optimal bin size decided by Freedman Diaconis Estimator shown in Fig. S1. The resulting sgFC values corresponding to those without direct SC link are mapped back to a 1000-by-1000 matrix and used for network analysis through the article.

### Diffusion map embedding

Diffusion map embedding is a nonlinear dimensionality reduction algorithm [11]. The algorithm seeks to project a set of embeddings into a lower-dimensional Euclidean space. Briefly, the similarity matrix among a set of points (in our case, the correlation matrix representing functional connectivity) is treated as a graph, and the goal of the procedure is to identify points that are proximal to one another on the graph. In other words, two points are close together if there are many relatively short paths connecting them. A diffusion operator, representing an ergodic Markov chain on the network, is formed by taking the normalized graph Laplacian of the matrix. The new coordinate space is described by the eigenvectors of the diffusion operator. We set the diffusion rate *α* = 1 and the variance of the Gaussian used in affinity computation *σ* = 1. The procedure was implemented using the Dimensionality Reduction Toolbox (https://lvdmaaten.github.io/drtoolbox/) [49].

## ACKNOWLEDGMENTS

We thank Justine Hansen, Vincent Bazinet, Golia Shafiei, Estefany Suarez, Andrea Luppi and Filip Milisav for their comments and suggestions on the manuscript. This research was undertaken thanks in part to funding from the Canada First Research Excellence Fund, awarded to McGill University for the Healthy Brains for Healthy Lives initiative. BM acknowledges support from the Natural Sciences and Engineering Research Council of Canada (NSERC Discovery Grant RGPIN #017-04265) and from the Canada Research Chairs Program.

---

**Figure S1.**
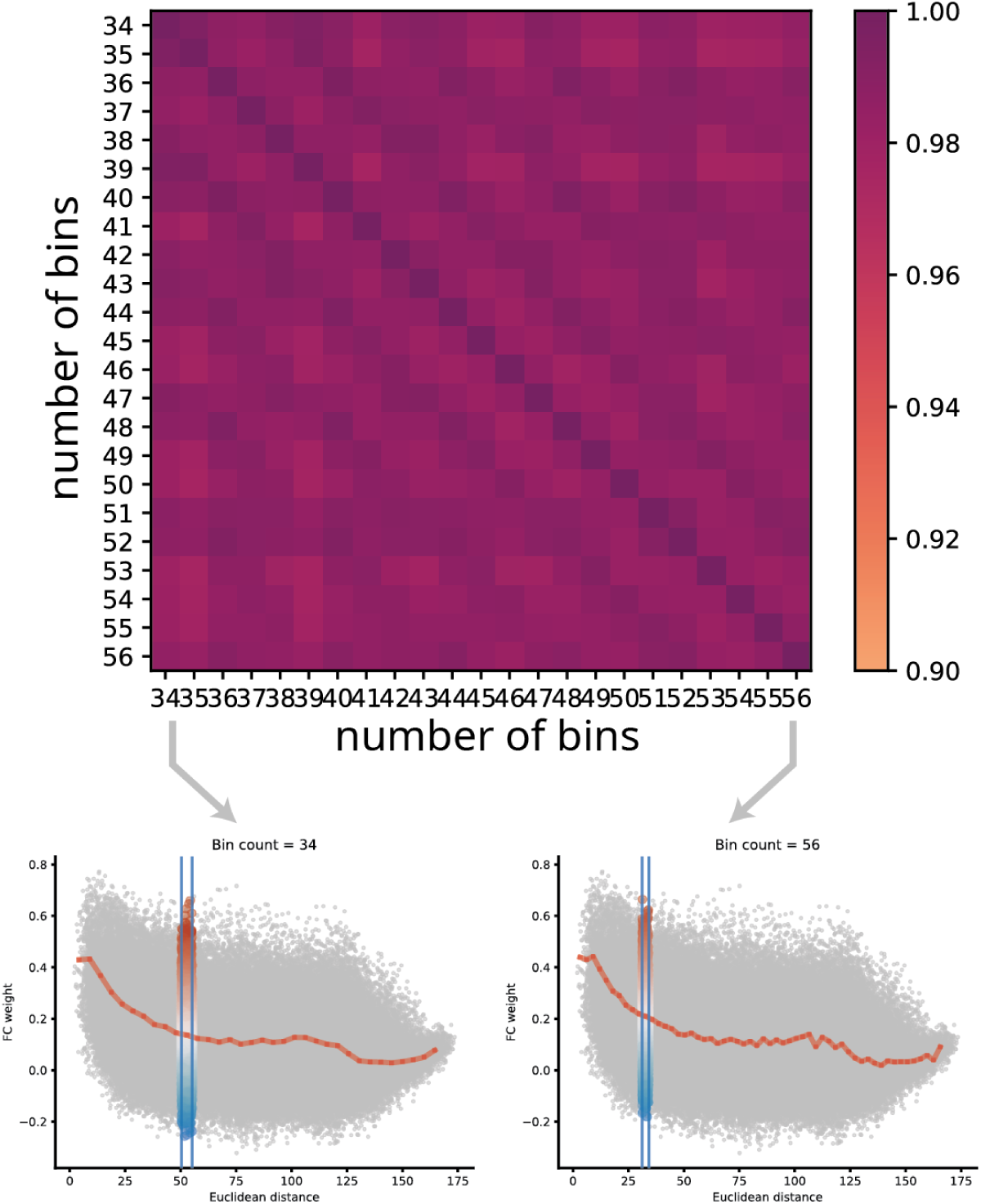
Influence of bin choice. Correlation matrix shown for sgFC values between choices of bin numbers. The number of bins are taken *±*25% centering the optimal bin size decided by Freedman Diaconis Estimator. The final sgFC values are averaged over the choices of bin numbers to get a smoothed robust representation.

**Figure S2.**
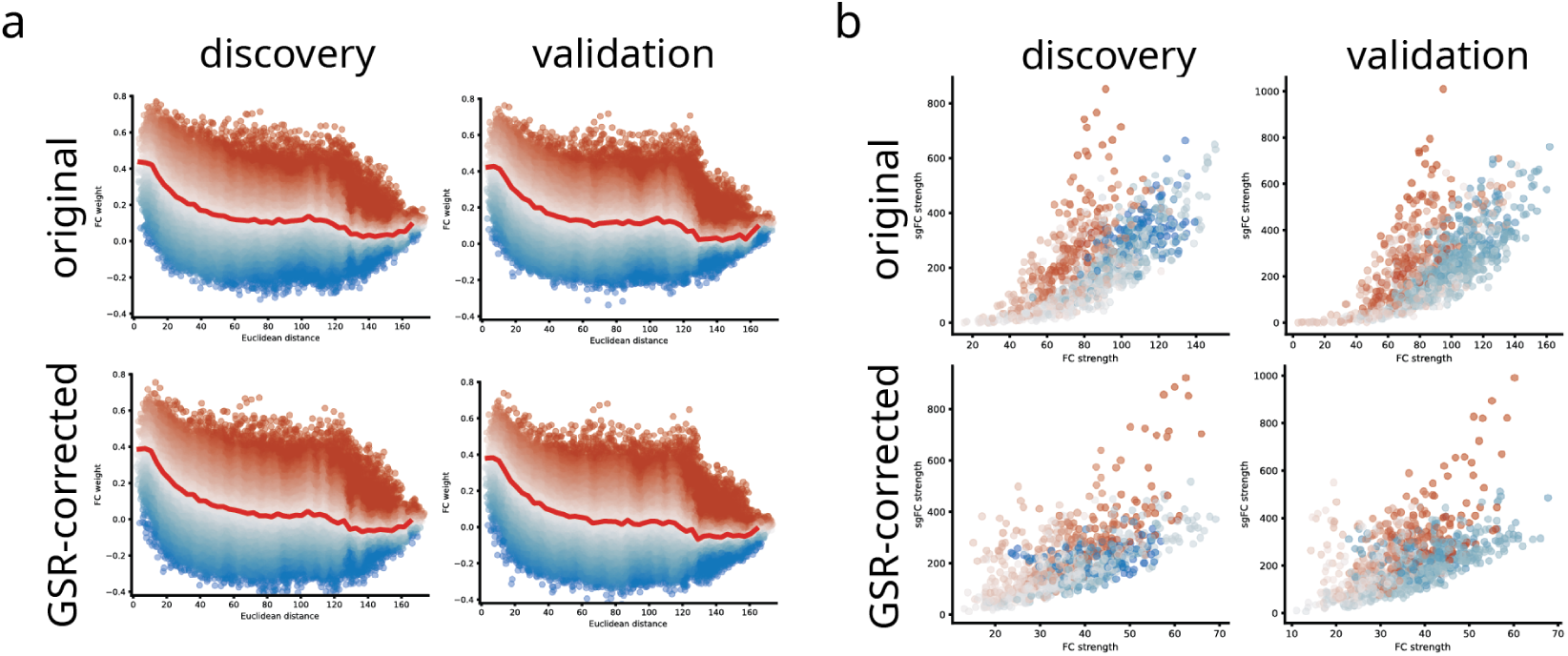
Results of control analyses. Reproduced Fig. 2d and Fig. 4b under control analyses settings. Correlations between the control cases for values in panel a (sgFC weight) and panel b (sgFC node strength) are shown in Fig. 5.

**Figure S3.**
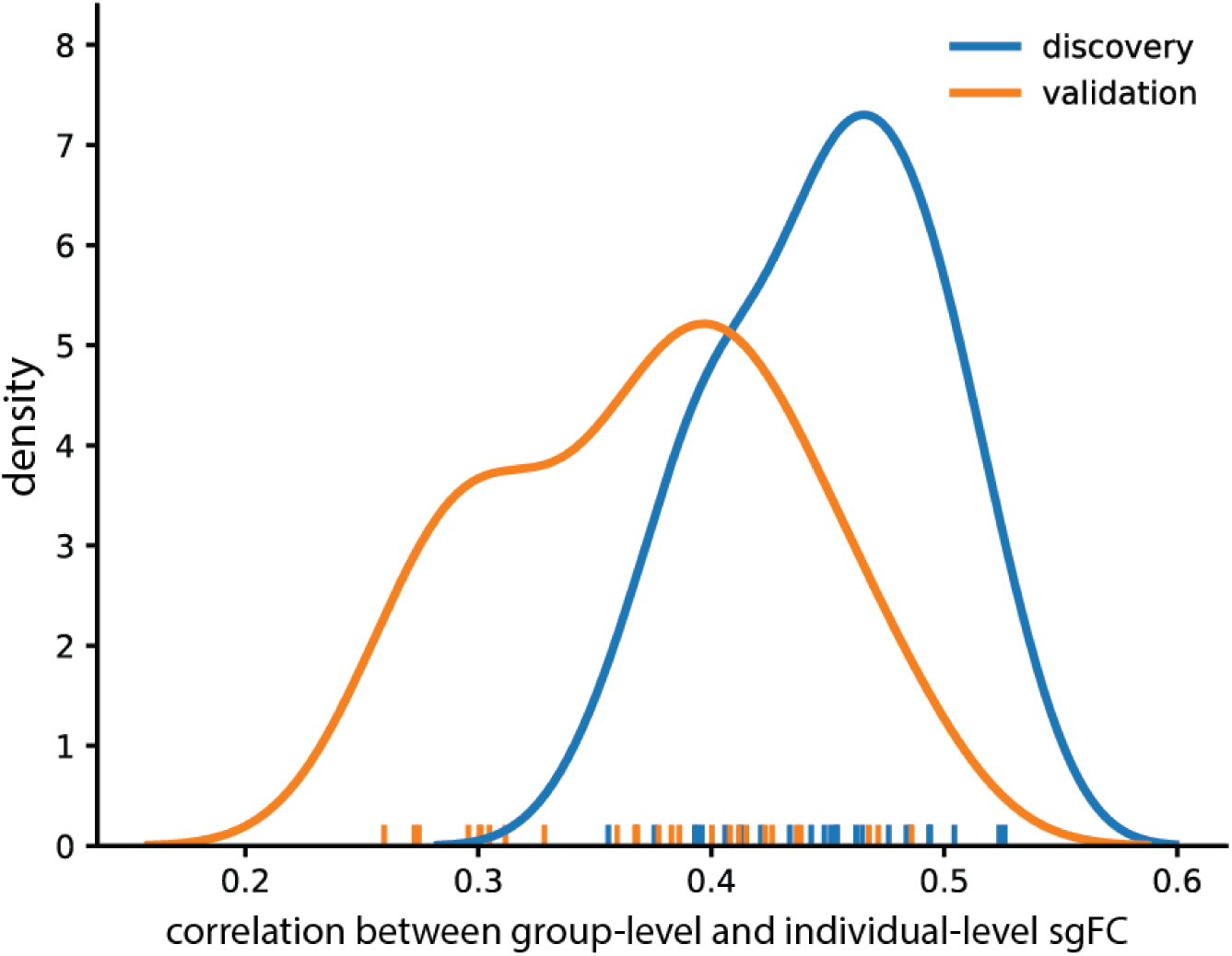
sgFC in individual participants. Distribution of correlation between group-level sgFC used in the main analysis and those for *N* = 66 participants. Group-level sgFC values for the *Discovery* dataset are correlated with sgFC generated from each participant in the *Discovery* and *Validation* dataset.

